# Variation and constraints in hybrid genome formation

**DOI:** 10.1101/107508

**Authors:** Anna Runemark, Cassandra N. Trier, Fabrice Eroukhmanoff, Jo S. Hermansen, Michael Matschiner, Mark Ravinet, Tore O. Elgvin, Glenn-Peter Sætre

**Author notes:** Corresponding author. (A.R.).

## Abstract

Recent genomic investigations have revealed hybridization to be an important source of variation, the working material of natural selection^1,2^. Hybridization can spur adaptive radiations^3^, transfer adaptive variation across species boundaries^4^, and generate species with novel niches^5^. Yet, the limits to viable hybrid genome formation are poorly understood. Here we investigated to what extent hybrid genomes are free to evolve or whether they are restricted to a specific combination of parental alleles by sequencing the genomes of four isolated island populations of the homoploid hybrid Italian sparrow *Passer italiae*^6,7^. Based on 61 Italian sparrow genomes from Crete, Corsica, Sicily and Malta, and 10 genomes of each of the parent species *P. domesticus* and *P. hispaniolensis,* we report that a variety of novel and fully functional hybrid genomic combinations have arisen on the different islands, with differentiation in candidate genes for beak shape and plumage colour. There are limits to successful genome fusion, however, as certain genomic regions are invariably inherited from the same parent species. These regions are overrepresented on the Z-chromosome and harbour candidate incompatibility loci, including DNA-repair and mito-nuclear genes; loci that may drive the general reduction of introgression on sex chromosomes^8^. Our findings demonstrate that hybridization is a potent process for generating novel variation, but variation is limited by DNA-repair and mito-nuclear genes, which play an important role in reproductive isolation and thus contribute to speciation.

---

Introgressive hybridization can transfer adaptive genetic variation across species boundaries^4^ and generate new species^2,4,5^. Although historically thought to be unimportant in animals, the genomic revolution has shown hybridization to be a pervasive evolutionary force, common in both plants and animals,^1,2^ and one that has even shaped the genome of our own species^8^. Portions of the genome vary in extent of introgression^4,9^, and homoploid hybrid genomes are predicted to have unequal parental contributions^10^. However, how hybridization, and subsequent recombination and selection mold genomes is not well understood. For instance, it is not known what determines the relative contribution of each parent and whether the genomic locations of introgression are subject to constraints or are largely stochastic. Moreover, genomic constraints in hybridizing taxa can be used to identify genes involved in reproductive isolation between species as they do not introgress^7^, and hence are important for our understanding of speciation and how biodiversity arises. One well-supported finding is reduced introgression on sex chromosomes, which is common in species where one sex is heterogametic^8,11^. To what extent this pattern is caused by specific types of genes on sex chromosomes is debated^12^, and knowledge of the genes causing incompatibilities and reproductive isolation is needed to resolve this.

To investigate whether certain genes must be inherited from a specific parent species to form a stable and functional hybrid genome and whether divergent genomes can arise from hybridization, genome-wide data from multiple independent hybrid populations are required. We studied isolated island populations of the homoploid hybrid Italian sparrow *Passer italiae*^6,7^ (Fig. 1a) to determine the constraints on, and variability of, genome regions and gene categories. The Italian sparrow genome is a mosaic of the genomes of its parent species, the house *P. domesticus* and Spanish *P.hispaniolensis*^13^ sparrows. The species is thought to have originated when house sparrows colonized the Mediterranean less than 10,000 years ago^6,14^. Morphologically divergent populations of Italian sparrows are found on the Mediterranean islands of Crete, Corsica, Sicily and Malta (Fig 1a; ^15^) and we used these isolated island populations to investigate genome-wide patterns of divergence and differentiation from the parent species and within the hybrid species (Supplementary Tables 1-2). We sequenced 10-21 Italian sparrow genomes from each island and 10 genomes from each of the parent species to 6-16X coverage as well as a tree sparrow (*P. montanus*) as an outgroup and aligned them to the recently *de novo* assembled house sparrow reference genome^13^.

**Figure 1.**
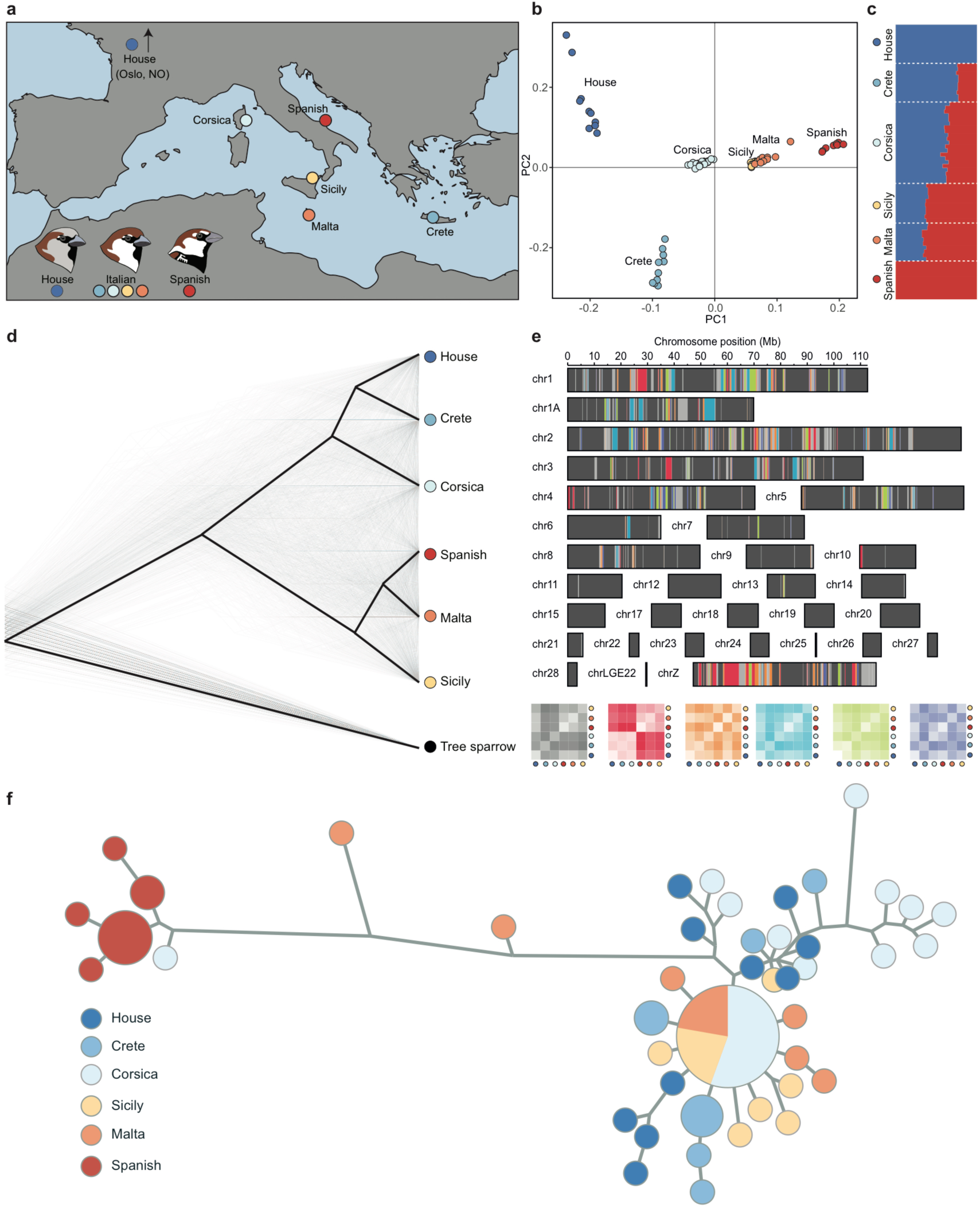
Population structuring of the focal taxa. **a**, Map showing the location of the island populations and the reference parent species populations, with examples of male plumage patterns. **b**, PCA of LD-pruned high quality SNP set with eigenvector 1 on the x-axis and eigenvector 2 on the y-axis. **c**, Population structuring based on Admixture analysis for house sparrow, Italian and Spanish sparrow populations. **d**, BEAST trees illustrating genome-wide variation in phylogenetic clustering between the taxa. **e**, SAGUARO plot illustrating the distribution of the six most common relatedness matrixes over the genome. **f**, Haplotype genealogy graph of mitochondrial sequences.

To determine if these Italian sparrow island populations were genomically differentiated, we first used a principal component analysis. Our results show strong support for differentiation among the hybrid island populations and from the parent species (Fig 1b; Supplementary Fig. 1; Supplementary Tables 3-4). The Italian sparrow populations differ in position along the axis of divergence of the parent species, with Crete and Corsica closer to house sparrow and Sicily and Malta closer to Spanish sparrow (Supplementary Table 5). To further investigate population differentiation, we assessed the most likely number of genetic clusters in the data, and the individual probability of belonging to these clusters using a structure analysis. We found support for two clusters (Supplementary Table 6) corresponding to the parent species and intermediate assignment probabilities for the Italian populations, with Crete and Corsica having the highest probabilities of clustering with house sparrow and Sicily and Malta the highest probabilities of clustering with Spanish sparrow (Fig. 1c). The assignment probabilities differed significantly among populations (ANOVA F_3,47_=736.54, *P*=2.2e-16; Supplementary Table 7), and across the genome (Supplementary Fig. 2). Moreover, in the most frequent phylogenetic tree topology, house sparrow, Crete and Corsica form one cluster and Spanish sparrow, Malta and Sicily the other, with Crete and Malta clustering most closely with parent species (Fig. 1d and Supplementary Fig. 3). The phylogenetic clustering varied spatially over the genome (Fig. 1e; Supplementary Table 8). We also found significantly higher Spanish sparrow introgression in the Sicily and Malta populations (Patterson’s D: ANOVA F_3,115_=22.52, *P*<0.001; Supplementary Table 9). The non-recombining mitochondrial DNA was similar to that of house sparrow for all Italian sparrows, with the exception of one Corsican individual having Spanish sparrow mitochondrial DNA, and two Maltese individuals appearing to have both house and Spanish sparrow mitochondria (Fig. 1f). This is consistent with heteroplasmy, as previously reported for mainland Italian sparrows^13^.

To address whether differentiation within a hybrid species is likely to result from positive selection or if differentiated areas mainly arise as a by-product of long-term linked background selection in low recombination areas^16^, we first investigated whether differentiation among Italian populations correlates with that between the parent species. We then tested whether variation in differentiation reflects variation in recombination rate, as has previously been shown in flycatchers^16^. Differentiation among the Italian sparrow populations was not strongly correlated with differentiation between the parents, (Supplementary Fig. 4). The standardized slope (beta) of the relationship between within-Italian differentiation and recombination rate was much shallower than that between the parent species differentiation and recombination rate (Supplementary Fig. 4; Tables S10-S11), and we therefore find no strong evidence for house sparrow recombination rate accounting for the genomic heterogeneity observed within this hybrid species. Sorting of parental variants, potentially due to Hill-Robertson effects^17^, may have contributed to this reduction in differentiation in low recombination regions compared to that of the parent species. Regions with high differentiation may instead reflect divergent selection, or, alternatively, the recombination landscape may be altered following hybridization and spur differentiation in other areas of the genome in the hybrid than in its parents.

To identify regions under divergent selection in the hybrid Italian sparrow we extracted the windows where the island populations were most divergent (measured as relative similarity to the parent species in terms of *F*_ST_). Among the genes found in the 1% windows that are most divergent between the island populations, 43 different gene ontology (GO) categories were overrepresented relative to the rest of the genome (Supplementary Table 12). Among these categories were genes related to neuron function, including nervous system development, transmission of nerve impulse, and synaptic transmission (Supplementary Table 12), suggesting that these categories of genes have been under divergent selection. Interestingly, genes related to neuron function have also been targets of recent positive selection in great tits^18^. The outliers also included FGF10, a gene explaining beak shape divergence between Darwin’s finches^19^. Sicily and Crete were strongly divergent at this beak shape candidate gene and the Italian sparrow has previously been shown to exhibit adaptive beak shape divergence^20^ (Fig. 2a-c). A gene involved in feather development^21^ and melanogenesis^22^, wnt7A, was also a highly differentiated outliers among the otherwise genetically very similar (mean *F*_ST_=0.016) but plumage-wise divergent Sicilian and Maltese populations^15^ (Fig. 2d-f and Supplementary Table 12). These two examples underscore that repeated hybridization between the same parental species can generate locally adapted populations through reshuffling of parental alleles at biologically important loci.

**Figure 2.**
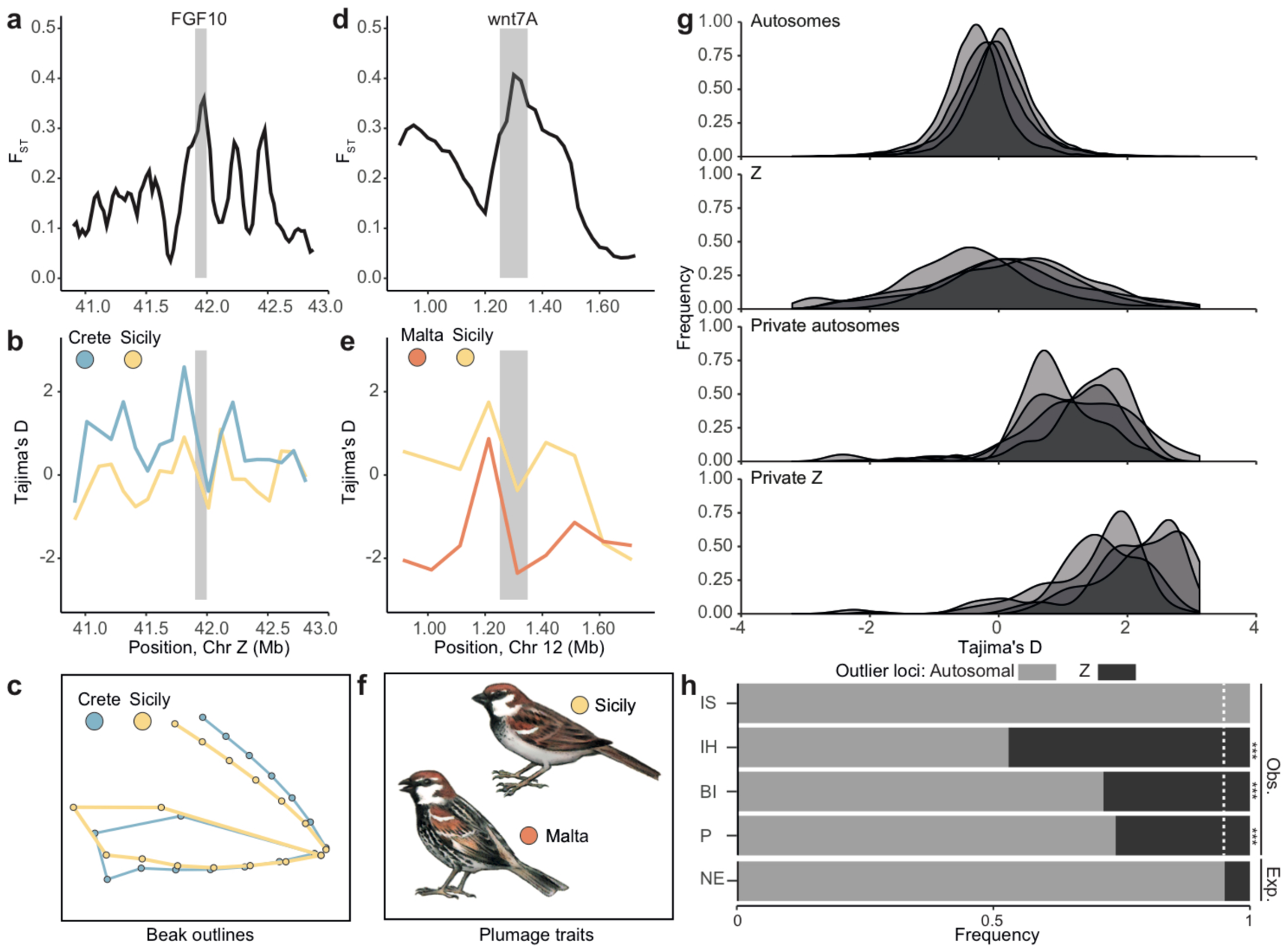
Local adaptation, private variation, and strong selection on the Z-chromosome. **a**, The beak shape candidate gene FGF10 is differentiated between Italian sparrows on Crete and Sicily. **b**, Tajima’s D for the FGF10 region. **c**, Beak shape differences between Cretan and Sicilian sparrows. **d**, The plumage candidate gene wnt7a is differentiated between genetically similar sparrow populations on Sicily and Malta. **e**, Tajima’s D for the wnt7a region. **f**, Schematic illustration of plumage differences between the populations. **g**, Strength of selection on the autosomes, the Z-chromosome and on private outlier loci where the Italian sparrow is differentiated from both parent species. **h**, Distribution of outlier genes between the Z-chromosome and the autosomes. IS denotes invariably Spanish, IH invariably house, BI between islands, P private and NE neutral expectations.

To identify areas of unique Italian sparrow evolution, we targeted regions in which the Italian sparrow populations have diverged from both parents by extracting the 1% of windows exhibiting the largest differences in *F*_ST_ between each Italian/parent comparison, only keeping windows overlapping between hybrid/parent in both comparisons. We found 12 overrepresented GO categories (Supplementary Table 13) including circadian rhythm, entrainment of circadian clock and rhythmic process: these genes showed strong signals of stabilizing selection (Fig. 2g), and elevated linkage disequilibrium (Supplementary Table 14). These results illustrate how population specific selection in concert with the parental mosaic is able to form unique features in the genomes of hybrid populations.

The level of differentiation between the Italian sparrow and the parent species is elevated on the Z-chromosome compared to autosomes (Paired t-test; t_3_=-8.40; *P*=0.004; Supplementary Fig. 5-6). This is expected based on the lower effective population size of this sex chromosome and hence elevated rates of genetic drift^23^. However, increased differentiation on the Z-chromosome is also expected from the faster X(Z) effect of elevated rates of adaptive evolution on the macro sex chromosome due to hemizygous exposure^24^. Outlier loci are strongly overrepresented on the Z-chromosome (Fig. 2h; all *P’s*<0.001; Supplementary Table 15), except for the outliers in the category where Italian populations invariably had inherited Spanish sparrow alleles, none of which resided on the Z. Interestingly, Tajima’s D estimates for the Z-chromosome have significantly higher variance than those for the autosomes (Fig 2c; Repeated measures ANOVA F_1,3_=56.94; *P*=0.005; Supplementary Fig. 7-8) and dn/ds was higher on the Z-chromosome compared to autosomes (goodness of fit *P*<0.001 for fixed differences against both house and Spanish sparrows; Supplementary Table 16), supporting a role for selection driving strong Z-chromosome divergence.

Across taxa with heteromorphic sex chromosomes, introgression on sex chromosomes is reduced^8,11^. To detect loci potentially important in causing such reduction in introgression on the sex chromosomes, we identified regions invariably inherited from a specific parent across all populations. We summed the *F*_ST_ against house sparrow across island populations and subtracted the summed *F*_ST_ against Spanish sparrow before extracting the extremes at both ends of the distribution (the 2% of the windows with squared values most diverged from 0; Supplementary Table 17). We found strong evidence for genes invariably inherited from house sparrow, especially on the Z-chromosome. DNA damage stimuli were significantly overrepresented among these genes (*P*_DNArepair_=0.026). There were 11 mitonuclear loci and although these were not generally overrepresented (*P*_mitonuclear_=0.11), 7 were found in the areas on the Z-chromosome strongly constrained to house sparrow inheritance, including the previously identified candidate incompatibility gene HSDL2^7^ (Supplementary Table 18; Fig. 3). There were also 6 DNA mismatch-repair genes on the Z-chromosome, among them the candidate incompatibility gene GTF2H2^7^ which is involved in nucleotide excision repair (Fig. 3). This suggests that hybrid genome formation is restricted to uniparental inheritance for these gene classes.

**Figure 3.**
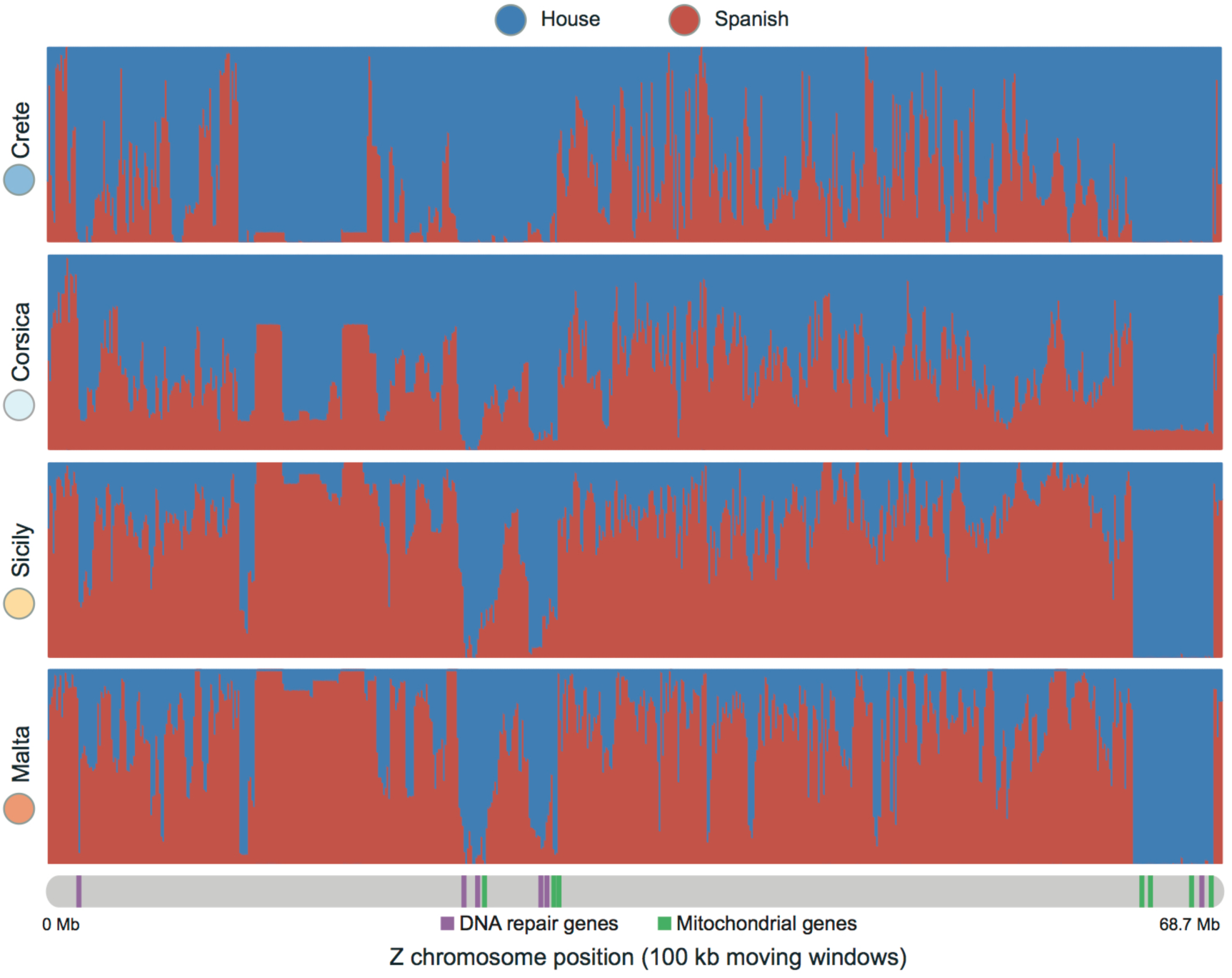
Parental similarity across the Z-chromosome. Sliding window ADMIXTURE analysis of probability of house sparrow (blue) or Spanish sparrow (red) inheritance over the Z-chromosome for the four Italian sparrow populations. Areas highly constrained to house sparrow inheritance harbour a significantly higher proportion DNA repair genes (green lines) and many mito-nuclear (purple lines) genes.

Whereas mito-nuclear genes are known as drivers of reproductive isolation^7^ and are expected to be under selection to interact with the frequent house sparrow-like mitochondria, the role for DNA repair genes is less established. However, reduced DNA repair functioning^25^ has been found in *Xiphophorus* fish hybrids, and the mismatch repair systems have been shown to contribute to meiotic sterility and cause incompatibilities in yeast^26^. Hence, multigenic DNA repair pathways may need parent specific inheritance to function. As most of these outlier genes were located on the Z-chromosome, they may contribute to the pattern of reduced introgression on sex chromosomes^7^.

Our comparison of isolated homoploid hybrid populations formed from the same parent species combination reveals that hybridization can produce diverged genomes with a range of different proportions of parental contribution. More outlier genes exhibited invariable inheritance from house sparrows than from Spanish sparrows. This suggests that parts of the genome must be inherited exclusively from one of the parent species, while the rest of the genome may vary with respect to parent species inheritance. Purging of Dobzhansky-Muller incompatibilities^27^ has been suggested to be important for shaping hybrid genomes. We find that house sparrow DNA-repair genes and mito-nuclear genes are necessary for “escaping the mass of unfit recombinants”^28^, most likely due to epistatic interactions. Genes invariably inherited from the Spanish sparrow are fewer but include a candidate pigmentation gene WNT4^29^ and a gene involved in vision, OLFML2B^30^, hence affecting external phenotype rather than genome function. Hence, both genome and organismal function can constrain hybrid genome formation, and the relative importance of the two may vary both quantitatively and qualitatively with the parent species.

Our data suggests that hybridization is a more potent force for creating novel variation than previously recognised, as many different combinations of the parental genomes can arise in hybrids and allow for adaptive divergence between isolated populations of the hybrid species. Importantly, we show that the variation is limited for DNA-repair and mitochondrial genes. These may contribute to the general pattern of reduced introgression on sex chromosomes, and are candidate loci for reproductive loci that may be important in speciation.

## Methods

### Field sampling

Italian sparrows were caught on Crete, Corsica and Sicily in 2013 and on Malta in 2014, while Spanish sparrows were caught in Lesina, Italy, 2008. House sparrows were caught in Northern Norway between 2007 and 2013, and the outgroup tree sparrow was caught in Sicily during 2008 (Supplementary Table 1). All sparrows were caught using mist nets, and blood was sampled from the brachial vein and immediately stored in Queens lysis buffer. Sparrows were released immediately after blood sampling to minimize stress. All permits were obtained from appropriate authorities prior to sampling.

### Whole genome resequencing, data processing and analysis

### DNA extraction and sequencing

DNA was extracted from blood samples stored in Queens lysis buffer using Qiagen DNeasy Blood and Tissue Kits (*Qiagen Corp*., Valencia, CA), and stored in Qiagen Elution Buffer (*Qiagen Corp*., Valencia, CA). Whole genome re-sequencing was performed with Illumina sequencing technology. An Illumina TruSeq gDNA 180 bp library was created and sequenced on the Illumina HiSeq 2000 platform with 100 bp read length and three individuals per lane for the parent species, the tree sparrow and the Italian sparrows from Malta; sparrows from Crete, Corsica and Sicily were sequenced with four individuals per lane. All re-sequencing was performed by Genome Quebec at McGill University (Montreal, Canada) (http://www.genomequebec.com/en/home.html). Raw data have been deposited at the NCBI Sequence Read Archive under (Accessions YYYY).

### Variant calling and filtering

All raw sequence reads were mapped to a repeat-masked version of the house sparrow genome^13^ using BWA 0.7.8^31^ with the mem, -M and -R options. A sorted BAM file was produced using SAMTOOLS version 1.0^32^ using view with the -b, -U and -s options and a pipe to the sort command. Duplicates were identified and filtered out using MARKDUPLICATES from PICARD-TOOLS version 1.107 (http://broadinstitute.github.io/picard/) using the options validation stringency = lenient, assume sorted = true and index = true. Indels were identified using RealignerTargetCreator and local realignments around these were performed with IndelRealigner. Standard settings were used for both these tools which are components of GATK 3.3.0^33,34^. Final sequencing coverage of these final BAM files (excluding duplicates) was 8x per individual (min. 5.99x, max. 15.8x; Supplementary Table 2). Variants were then called using the GATK HaplotypeCaller. First, HaplotypeCaller was run separately for each individual to create single-sample gVCFs using the –emitRefConfidence GVCF, –variant_index_type LINEAR and –variant_index_parameter 128000 options, and then GATK GenotypeGVCFs tool was run using standard settings to achieve joint genotyping. Two different versions of the VCF file were created one with variable sites only (49 237 560 SNPs), and one where all sites were called as specified by the –includeNonVariantSites option (1 040 518 317 SNPs).

For both VCF files, indels were first filtered out using VCFtools version 0.1.12b^35^, and hard filtering according to the Broad Institute’s recommendations was performed with bcftools-1.2^32^ in order to filter out sequencing artefacts. This included requiring a QualByDepth of at least 2.0 and a FisherStrand phred-scaled p-value of less than 60.0, based on Fisher’s Exact Test to detect strand bias in the reads, which may be indicative of false positive calls. Hard filtering also required a RMSMappingQuality of at least 40.0 to ensure high mapping quality of the reads across all samples, a MappingQualityRankSumTest value of −12.5 to exclude reads where the alternate alleles has a lower mapping quality than reads with the reference allele, and finally a ReadPosRankSumTest value of less than -8.0 was required to ensure that reads with the alternate allele were not shorter those with the reference allele, potentially indicating sequencing artefacts. In addition, we filtered out sites with mean number of reads per individual lower than 3, with a genotype quality lower than 20 and with a mapping quality lower than 20 using VCFtools version 0.1.12b^35^. To avoid paralogs, we also excluded sites with read depths above 5 times the variance of coverage. This left a total of 38 341 426 sites for further analysis.

### PCA

Principal component analysis was performed using ANGSD^36^ version 0.911 and ngsTools version 1.0.1^37^. This pipeline was chosen because it does not rely on genotype calls but instead takes allele frequency likelihoods and genotype probabilities into account^38^. We first estimated genotype probabilities from BAM files with ANGSD, including bi-allelic sites only and allowing a minimum mapping and site quality of 20 (Phred score) and a minimum coverage of 30x across all individuals. PCA was then performed using ngsTools on genotype probabilities. Allele frequencies were normalized and genotypes were not explicitly called, as specified by setting the options -norm 0 and -call 0 options. Eigenvalues for each PC were then estimated from the covariance matrix produced by ngsTools and data was plotted using a custom R script. We used the broken stick criteria to assess which PC axes were biologically informative from a simple scree plot and then extracted the covariance from these axes (Supplementary Table 3). The analysis was performed on three datasets: all sites, sites from the Z chromosome only, and autosomal sites only.

### Population genomic analysis

Population genetic parameters were estimated for non-overlapping 100 kb windows along the genome, as this window size was larger than the distance of LD-decay^13^. Population genetic inference was based on genotype likelihoods whenever possible. ANGSD^36^ version 0.911 was used to estimate allele frequency likelihoods and to obtain a maximum likelihood estimate of the unfolded site frequency spectrum (SFS) for estimation of Tajima’s D. Nucleotide diversity was estimated by dividing the pairwise Theta (population scaled mutation rate) estimates by the number of variable sites per window. The ancestral sequence was reconstructed using genotypes from the outgroup tree sparrow. A Fasta file of the tree sparrow genome was obtained by using the -doFasta 2 command with the -GL 1 -doCounts 1, -setMinDepth 3 and -setMaxDepth 65 options in ANGSD^36^ version 0.911. Here, BAM files from eight additional tree sparrows, sequenced to 8-12x depth and processed as described for the other samples above, were used. Genetic differentiation (*F*_ST_) based on genotype likelihood was estimated based on the two-dimensional SFS using ngsTools^38^. All these analyses were performed on the final BAM files. Sequence divergence (*d*_xy_) was calculated from the VCF file with all positions called using the script developed in^39^ (https://github.com/johnomics/Martin_Davey_Jiggins_evaluating_introgression_statistics/blob/master/egglib_sliding_windows.py; version August 2014).

### Admixture analysis

Genetic admixture was estimated using ADMIXTURE^40^ version 0.911. The VCF-file was converted to plink’s PED format using VCFtools version 0.1.12b^35^ and plink version 1.07^41^. Log likelihood values for K, the number of genetic clusters in the datasets, between K=1 and K=8 were estimated (Supplementary Table 4), and admixture analyses were run for the most appropriate value of K. Analyses were first run for a LD-pruned whole genome dataset (sites within a 50 SNP stepping window with a correlation coefficient higher than 0.1 were omitted; pruning of the BED file was performed with plink version 1.07 using the –indep-pairwise command; this left 438,443 sites for analysis). Sliding window analyses with 100 kb windows were then carried out to investigate variability in probability of parental inheritance across the genome. This analysis was performed individually for each island population together with the two parent species. The VCF-file was then split into individual VCF files per 100 kb using the –L option in GATK 3.3.0^33,34^. Individual ADMIXTURE analyses for K=2 were then performed for each 100 kb BED file. A mean estimated cluster assignment probability for all individuals per population was computed for each analysis using a custom python script.

### Phylogenetic analyses

To investigate whether phylogenetic relationships varied across the genome due to introgression or incomplete lineage sorting, a machine-learning approach implemented in SAGUARO version 0.1^42^ was used to identify genomic regions characterized by distinct similarity matrices. To focus on breakpoints at which recombination led to changes in the topology of populations or species, rather than topological changes within populations, one representative high-coverage individual per Italian sparrow population or sparrow species was used in these analyses. These were the house sparrow 8L19786, the Spanish sparrow Lesina_280 and the Italian sparrows C081 (Crete), K035 (Corsica), S059 (Sicily) and M036 (Malta). The program VCF2HMMFeature (included in the SAGUARO package) was used to convert the VCF-file to the HMMFeature format required by SAGUARO, and SAGUARO was run using default parameters. Analysis was performed jointly for all chromosomes, as the same similarity matrices are expected to occur on multiple chromosomes. A total of 797 contiguous regions of up to 54,398 bp were identified and assigned to one out of 41 similarity matrices, of which the most common similarity matrix characterized 79.08% of the genome. Similarity matrices represented by more than 2% of the genome are depicted in Figure 1.

The genomic regions identified by SAGUARO were subsequently used to infer differences in phylogenetic relationships more thoroughly with the Bayesian software BEAST version 2.2.0^43^. To this end, chromosome-length alignments were first phased using SHAPE-IT version 2^44^. To improve phasing, this analysis was conducted with a subset consisting of the six individuals with the highest coverage per population (24 individuals in total) rather than just the six individuals used for SAGUARO analyses, however, the six focal individuals were extracted from the alignments following phasing. The phased chromosome-length alignments were then used to extract 38,964 non-overlapping blocks of 25,000 bp from the 797 contiguous regions identified with SAGUARO. For each of the 38,964 blocks, one of the two phased sequences per individual was excluded at random, so that each alignment contained a single sequence per population or species. To identify alignments particularly suitable for Bayesian phylogenetic analysis, we quantified, for each alignment, the proportion of missing data, the number of parsimony-informative sites, the proportion of heterozygous sites, the mean bootstrap support of maximum-likelihood trees generated with RAXML version 8.2.4^45^, and the probability that the alignment is free of recombination determined with the Phi test^46^. We assumed that alignments with a low proportion of heterozygous sites are less likely to contain paralogous sequences, and that alignments with many parsimony-informative sites and high mean bootstrap support contain strong phylogenetic signal. Thus, alignments were selected according to the following “relaxed” and “strict” filters: a proportion of missing data below 0.2 (relaxed) or 0.1 (strict), at least 75 (relaxed) or 100 (strict) parsimony-informative sites, a proportion of heterozygous sites below 0.005 (relaxed) or 0.0025 (strict), a mean bootstrap support of at least 90 (both relaxed and strict), and a Phi test *p*-value above 0.005 (relaxed) or 0.01 (strict). A total of 1,234 and 116 alignments were selected with these relaxed and strict filters, respectively. To include an outgroup for phylogenetic analyses with BEAST, consensus sequences of tree sparrow reads from the Naxos1 individual mapped to the house sparrow reference genome^13^ were added to each selected alignment. To avoid bias towards the reference, missing data were not replaced by the reference alleles. The phylogeny of each alignment was then inferred with BEAST, using the GTR model of sequence evolution with estimated base frequencies, a Yule tree prior^47^, and 50 million Markov chain Monte Carlo iterations. The ingroup, combining house sparrow, Spanish sparrow, and the representatives of Italian sparrow populations was constrained to be monophyletic. In the absence of a reliable absolute time line for sparrow divergences, the time of divergence of ingroup and outgroup was fixed at 1 time unit, so that all divergence ages within the ingroup are estimated relative to this initial split. Convergence of all MCMC chains was confirmed by effective samples sizes greater than 500 for all model parameters.

### Introgression

Presence of introgression was estimated using Patterson’s D^48,49^ calculations, using the scripts provided in^39^. ABBA-BABA estimates were calculated using a minimum coverage of 3, a 100 kb window size and 1000 informative sites using the egglib_sliding_windows.py script. The test was set up to estimate Spanish sparrow introgression into a house sparrow background, with tree sparrow as an outgroup.

### Mitochondrial DNA

Mitochondrial DNA gvcfs were called separately with haploid settings using the -ploidy argument in HaplotypeCaller, jointly genotyped, and filtered as described above using GATK 3.3.0^33,34^. Fitchi version 1.1.4 ^50^ was used to reconstruct a haplotype genealogy based on Fitch distances.

### Recombination rate and common differentiation

Genome-wide recombination rates were estimated using a house sparrow linkage map. As the recombination map was produced using SNP chip data, recombination distance estimates were first averaged using a sliding window approach and then a loess fit of mean recombination rate against physical distance was performed in order to interpolate fine scale variation across non-overlapping 100 kb windows. Since recombination data were not available for the Z chromosome, this interpolation was performed on autosomes only.

To test whether there was a relationship between recombination rate variation and relative genomic differentiation, either *F*_ST_ from 100 kb non-overlapping windows, which is a direct measure of relative differentiation, or the common differentiation axis – i.e. shared differentiation amongst groups of populations – were used. The latter was calculated by performing PCA on multiple pairwise comparisons of differentiation featuring the same focal species following^51^. Common differentiation was estimated amongst 1) all Italian populations, 2) between all Italian populations and the house sparrow and 3) between all Italian populations and the Spanish sparrow. In each case, all pairwise *F*_ST_ comparisons including these species were included, and the first principal component extracted.

### Outlier gene analysis

Disparities in *F*_ST_ values between lineages were used to identify genomic regions in which the Italian sparrow populations display elevated divergence from either or both of their parents or other Italian sparrow populations. This approach is reminiscent to population branch statistics. Three categories of outliers were of interest: Between island outliers, where island populations differed strongly in parental resemblance, were selected as these are informative of how Italian sparrow populations are differentially adapted. Private outliers, windows in which Italian sparrows are diverged from both parent species, show where unique adaptation is putatively strongly selected for. Finally, portions of the genome invariably inherited from the same parent species for all populations are informative of parts that are under strong selection to resemble a specific parent species and may reveal constraints on hybrid speciation.

#### Between island outliers

The 1% of 100 kb regions which differed most with respect to *F*_ST_ against the parental species between two islands were selected for all possible island-island combinations. All genes within or partly within these regions were then extracted. As historical effects such as ancestral polymorphism and selection prior to the parental split can be assumed to be constant across Italian populations, using the difference in *F*_ST_ against the same parent species will yield results which are not dependent on these factors. Furthermore, *F*_ST_ was not strongly dependent on recombination rate between Italian sparrow populations (Supplementary Fig. 4).

#### Private Outliers

Outliers in which Italian sparrows were differentiated from both parent species, hereafter private outliers, were extracted from the 1% of windows exhibiting the largest difference in *F*_ST_ between each hybrid/parent comparison, only keeping windows overlapping between both hybrid/parent comparisons. This was done separately for each Italian population, and all outliers detected across the populations were then merged for gene ontology analyses.

#### Outliers invariably resembling one parent species

To limit historical effects, due to for instance ancestral polymorphism and selection prior to the parental split, we used the 100 kb windows in which the Italian sparrow had the cumulative largest difference in *F*_ST_ value between one parent and the other.

This was achieved by summing the *F*_ST_ values between all Italian sparrow populations and house sparrow, and subtract the sum of the *F*_ST_ values between these populations and Spanish sparrow. As the resulting distribution was skewed, using a percentage at each tail would not have captured the biological pattern where house sparrow inheritance across populations was more common than Spanish sparrow inheritance across all populations. Therefore, we squared the summed values and extracted the 2% of the windows that deviated most strongly from 0 (Supplementary Fig. 8B), which yielded more invariably house sparrow like outliers than invariably Spanish sparrow like outliers.

For all outlier windows in each of the three categories above, annotated genes that resided completely or partially within them were extracted for separate gene ontology analysis. One analysis was performed on all private outliers identified across populations, one on all outliers between populations, including all combinations of populations, one on outliers that resembled house sparrow across all populations, and finally one on outliers that resembled Spanish sparrow across all populations. As only 14 outliers resembling Spanish sparrow across all populations were found, no significant GO-terms were found for this analysis. Therefore, we do not provide a table of significant terms for this analysis. These analyses were performed using GO-stat^52^, with a human reference base. We implemented standard settings for GO analyses, i.e. a values of 3 as the minimal length of considered GO paths and no merging of GOs if gene lists overlap. Mito-nuclear genes were identified using MITOMINER 4.0^53^ with a human reference database and standard settings. Overrepresentation of mitonuclear genes was subsequently tested using a Chi-square test. Corrections for multiple testing were performed with the Benajmini method.

### DN/DS analyses

To test if the Z chromosome was under stronger selection, synonymous and non-synonymous fixed differences within genes against each parent species for all autosomes and the Z chromosome were estimated for each Italian sparrow population. A goodness of fit test was applied to test if the number of nonsynonymous substitutions on the Z chromosome was higher than expected for each parent species. To this end, the R package PopGenome^54^ was used. The splitting, data command was used to extract genes and fixed sites were extracted. Synonymous and nonsynonymous sites were then identified using the options subsites="syn" and subsites="nonsyn", respectively.

### Linkage disequilibrium decay

To address whether linkage disequilibrium (LD) was higher and LD decayed more slowly in outlier windows than in randomly selected windows, plink version 1.90b3b^41^ was used. Using –const-fid –ld-window 1000 –ld-window-kb 100 –r2 and –ld-window-r2 0.0, linkage disequilibrium in the 100 kb outlier windows was estimated within each of the Italian populations. In addition, we randomly selected 1,000 100 kb windows spread across the chromosomes in proportion with chromosome size and estimated linkage disequilibrium for these in the same manner for comparison. A linear model was fitted for each outlier window, and intercept and slope were recorded and used in glm’s to test whether LD was higher and LD decay was slower in outlier windows than in randomly selected windows.

**Supplementary Information** is linked to the online version of the paper at www.nature.com/nature.

## Acknowledgements

We thank Maria Tesaker and BirdLife Malta for help with field work, Laura Piñeiro and Lena Bache-Mathiesen for providing morphological data, and Anna Nilsson for comments on the manuscript. This work was funded by a Swedish Research Council post doctoral grant and a Wenner-Gren Fellowship to A.R. and a Norwegian Science Foundation grant to G-P.S. and A.R.

## Author Contributions

A.R. conceived the study, carried out field work, lab work, designed analyses, analysed data and wrote the manuscript. C.N.T. helped design analyses, and provided example scripts for many analyses, F.E. carried out field work and the gene ontology analyses, J.S.H. carried out field work and the final touches in figure preparation, M.M. did the BEAST and Saguaro analyses and M.R. performed the recombination rate analyses and PCA. T.E. provided the house sparrow reference genome, and G.P.S. identified the study system, designed the sampling strategy and carried out field work. All co-authors commented on the manuscript.

Data is deposited at YYYY. Reprints and permissions information is available at www.nature.com/reprints. We declare no competing financial interests.

Correspondence and requests for materials should be addressed to.

